# Linking physiology to ecosystem function: how vulnerable are different functional groups to climate change?

**DOI:** 10.1101/2022.06.07.495062

**Authors:** Carmen R.B. da Silva, Julian E. Beaman, Jacob P. Youngblood, Vanessa Kellermann, Sarah E. Diamond

**Affiliations:** Department of Biology, Case Western Reserve University, Cleveland, Ohio, USA; School of Biological Sciences, Monash University, Clayton, VIC 3800, Australia; College of Science and Engineering, Flinders University, Bedford Park, SA 5000, Australia; School of Life Sciences, Arizona State University, Tempe, Arizona 85287, USA

## Abstract

1. The resilience of ecosystem function under global climate change is governed by individual species vulnerabilities and the functional groups they contribute to (e.g. decomposition, primary production, pollination, primary, secondary and tertiary consumption). Yet it remains unclear whether species that contribute to different functional groups, which underpin ecosystem function, differ in their vulnerability to climate change.
2. It is important to examine if functional group vulnerability differs across space (e.g. tropical vs temperate latitudes) to determine if some regions will be more vulnerable to loss of ecosystem function than others, and to examine whether localized effects of particular community compositions override global patterns of functional group vulnerability.
3. We used existing upper thermal limit data across a range of terrestrial species (N = 1,743) to calculate species warming margins (degrees distance between a species upper thermal limit and the maximum environmental temperature they inhabit), as a metric of climate change vulnerability, to determine whether species that comprise different functional groups exhibit differential vulnerability to climate change, and if vulnerability trends change across geographic space.
4. We found that primary producers had the broadest warming margins across the globe (μ = 21.85 °C) and that tertiary consumers had the narrowest warming margins (μ = 4.37 °C), where vulnerability tended to increase with trophic level.
5. Species that contribute towards primary production were more vulnerable in low-latitude than mid-latitude regions, but warming margins across all other functional groups did not differ across regions when evolutionary history was considered. However, when evolutionary history was excluded from the analyses (as closely related species often play similar functional roles within ecosystems demonstrating true variation in functional group warming margins) we found that pollinators are more vulnerable in mid-latitude regions and that primary producers are more vulnerable in low-latitude environments.
6. This study provides a critical first step in linking individual species vulnerabilities with whole ecosystem responses to climate change.

## Introduction

The functional roles species play in ecosystems such as decomposition, primary production, consumption, and pollination scale up to support ecosystem function (Box1) (Crowther et al., 2015; Enquist et al., 2003, 2015). As climates change, and species respond individually via shifts in geographic ranges, phenology, or changes in population abundances (increase or decrease), we will observe alterations in functional group (Box 1) interactions and species compositions, which are anticipated to have major effects on ecosystem function (Harvey et al., 2020; Oliver et al., 2015; Voigt et al., 2003). The robustness of ecosystem function to climate change will depend, in part, on the diversity of species that contribute to each functional role (functional redundancy) (García et al., 2018; Hisano et al., 2018; Loreau, 2000). With greater species diversity there is an increased likelihood that some species will be resilient to warming climates, decreasing the likelihood that functional roles within ecosystems will be lost to climate change. However, if the vulnerability of species that contribute towards different functional groups within ecosystems is non-equal, we might observe declines in ecosystem function at a more accelerated rate than we would expect based on individual species vulnerabilities to climate change (Thackeray et al., 2016; Thakur, 2020; Voigt et al., 2003).

#### Box 1. Glossary

##### Ecosystem function

fluxes of energy, nutrients and organic matter through an environment which in turn supports ecosystem productivity and stability.

##### Functional groups

a group of species that contribute towards a certain *functional role* within an ecosystem such as: primary production, consumption (primary, secondary or tertiary), pollination and decomposition. We use the term functional group rather than trophic level because pollination is not a trophic level. But, as pollinators play one of the most important roles in plant reproduction, and thus primary production, it is an important functional role to consider within ecosystems.

##### Warming margin

the distance between an organism’s upper thermal limit and their (average warmest month) maximum environmental temperature. Species vulnerabilities to climate change increase with decreasing warming tolerance.

Ecological hypotheses predict that species that contribute towards higher trophic levels should be more vulnerable to climate change than lower trophic level species (Thackeray et al., 2016; Voigt et al., 2003). This is because large and highly active species are expected to lose the ability to maintain their energetic requirements under hot conditions faster than smaller and less active species (Brown et al., 2004; Huey & Kingsolver, 2019; Vasseur & McCann, 2005; Voigt et al., 2003). In addition, species that contribute towards lower trophic levels have a greater ability to shift their phenologies with climate change than higher trophic level species, potentially buffering them from warming temperatures (Thackeray et al., 2016). However, it remains unclear whether species that contribute towards different functional groups have different upper thermal limits, and thus different physiological vulnerabilities to climate change. Understanding how species thermal tolerances vary across the functional groups they contribute to can provide key information on both functional group vulnerability ranking and the degrees of climate warming until each functional role will be lost within ecosystems.

Species upper thermal limits can be estimated via tolerance assays (ramping or static) or thermal performance curves (Angilletta Jr, 2009; Bennett et al., 2021; Diamond et al., 2012; Kellermann et al., 2012; Sunday et al., 2014). Upper thermal limits can then be compared to current or future environmental temperature yielding a ‘warming margin’, or the temperature difference between a species upper thermal limit and the maximum environmental temperature they experience. Owing to their composite nature, warming margins can vary according to properties of the environment and properties of the organism (Kellermann et al., 2012; Sunday et al., 2014). In aggregate, warming margins identify the species or populations that inhabit environments close to their physiological capacities and thus are most vulnerable to climatic change (Kellermann et al., 2012; Sunday et al., 2014). Comparisons of species warming margins across spatial scales allows assessments of which geographic regions, such as temperate vs tropical, are most likely to lose species or populations to climate change. Whether tropical or temperate species are more vulnerable to climate change has generated great debate over the past 20 years, as species that inhabit tropical environments sit closer to their upper thermal limits on average, but temperate environments often experience higher extreme summer temperatures which have the potential to adversely impact populations (Deutsch et al., 2008; Helmuth et al., 2002; Kellermann et al., 2012; Kingsolver et al., 2013; Peralta-Maraver & Rezende, 2021; Sunday et al., 2014). While warming margins have been compared across species and latitude in the past, it is not known if species that contribute towards certain functional roles are more vulnerable than others, and if trends in functional group vulnerability changes with variation in species composition across ecosystems. Loss of certain functional roles in ecosystems could impose trophic cascades under climate change, potentially reducing ecosystem function (Thakur, 2020; Voigt et al., 2003).

In order to test if functional groups have differential physiological vulnerabilities to climate change, we leveraged large, compiled datasets of terrestrial species thermal tolerance and categorised each species into their principal functional group across the globe (tertiary, secondary, and primary consumption, pollination, primary production, decomposition). To compare the vulnerability of species that contribute to different functional groups, we calculated each species’ warming margin, and assessed how warming margins varied across functional groups. To provide nuance to the debate on whether tropical or temperate species are the most vulnerable to climate change, we examined how functional group vulnerability changes across low- and mid-latitude regions. Furthermore, to explore if trends in functional group vulnerability are maintained across geographic scale, and with changes in species composition across ecosystems, we compared global vulnerability trends with regional trends, and examined functional group vulnerability trends in a local scale case study, south-eastern Australia (Supplementary Analysis 1).

We appreciate that thermal limits and warming margins are estimated with caveats (e.g. variation in experimental methodologies), as highlighted in a number of manuscripts (Allen et al., 2016; Clusella-Trullas et al., 2021; Diamond & Yilmaz, 2018; Hoffmann et al., 2013). Here, we assume that thermal limits and their associated warming margins are a reasonable proxy for species vulnerabilities to climate change. This assumption is supported by findings that species’ upper thermal limits are correlated with warming-induced range shifts and extinctions (Comte et al., 2014; Diamond et al., 2012; Pinsky et al., 2019; Sinervo et al., 2010). We have also assumed that estimates of warming margins are static and unable to shift via evolution or plasticity. We appreciate that this is unlikely to be the case, but sufficient data do not currently exist to factor evolutionary potential or plasticity of thermal tolerance into our models. Nevertheless, the extent to which either plasticity or selection on heritable genetic variation will shift upper thermal limits is expected to be small (Gunderson & Stillman, 2015; Kellermann & van Heerwaarden, 2019).

Here we examine three main research questions. 1) Do species that contribute towards different functional groups vary in their vulnerability to climate change at a global level? 2) Does functional group vulnerability differ across tropical and temperate regions? 3) Are trends in functional group vulnerability robust to geographic scale and community composition? Finally, we discuss the limitations of our dataset and analyses, and discuss research priorities for the future.

## Materials and Methods

We compared upper thermal limit data of terrestrial species from the large databases GlobTherm (Bennett et al., 2018) and Lancaster & Humphreys (2020). Pollinators, decomposers and tertiary consumers were under-represented within these databases, so we filled in thermal tolerance data gaps by searching a set of specific search criteria for each functional group on Web of Science (https://www.webofscience.com/wos/woscc/basic-search) (see Supplementary Material Section 1 for criteria lists). In total, we compared 2,150 upper thermal limit data points across 6 functional groups (decomposers, primary producers, pollinators, primary consumers, secondary consumers, and tertiary consumers), and 1,743 species (Figure 1; see list of taxonomic classes within each functional group in global and regional samples in Supplementary Information 2). Some primary producer species had upper thermal limit data for more than one location and thus occurred in the dataset more than once. A full systematic review of all terrestrial species’ upper thermal limits was not conducted as this was accomplished already by Globtherm (Bennett et al., 2018) and Lancaster & Humphreys (2020). Missing taxonomic information for species was extracted from the National Centre for Biotechnology Information database (https://www.ncbi.nlm.nih.gov/taxonomy) using the Taxize package (Chamberlain & Szöcs, 2013) in the statistical program R version 4.1.0 (R Development Core Team, 2019). These large databases include upper thermal limit estimates obtained using a variety of methodologies such as dynamic, static and upper edge of the thermal neutral zone (TNZ) which we acknowledge could influence upper thermal limit estimations (Allen et al., 2016; Diamond & Yilmaz, 2018). Accordingly, we included methodology category (dynamic, static or TNZ) as a random factor within our analysis (summary of the methodologies used per functional group is shown in Supplementary Table 1).

**Figure 1.**
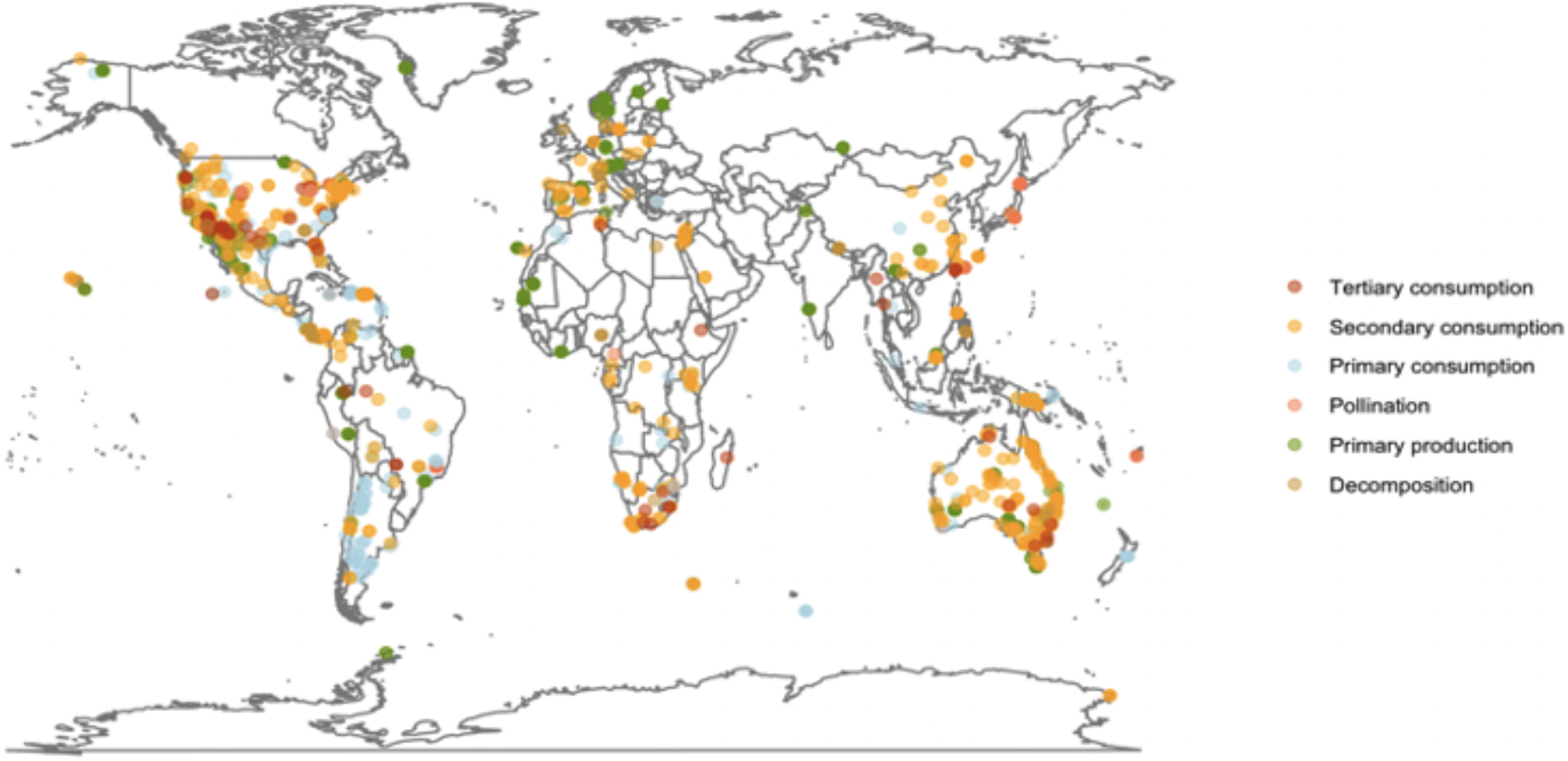
Collection locations of terrestrial species upper thermal limit data across the globe (N = 2,150). Points are coloured by the functional group that species contribute to.

We categorised species into functional group trophic level categories as per Reichle (2019). For simplicity, species were categorised into functional groups based on their predominant adult life stage energy sources to the family level, i.e. autotrophs were labelled primary producers and species in families that predominantly eat primary producers were considered primary consumers. Species in families that predominantly eat primary consumers were considered secondary consumers and those that eat secondary consumers were considered tertiary consumers. Species in families with known decomposition and pollination roles were also categorised accordingly. However, some species were considered both primary consumers and pollinators and thus appear twice in the dataset, and one species of lichen (*Lobaria pulmonaria)* is considered both a decomposer and a primary producer. We then assessed how functional group vulnerability in the above categories differs across a global, regional (low vs mid latitudes) and local (south-eastern Australian forests - Supplementary Analysis 1) scales.

### Analysis

Analyses were performed in the statistical program R version 4.1.0. Warming margins for each species were calculated by subtracting the mean maximum environmental temperature (BioClim variable 5) (Fick & Hijmans, 2017) at each species collection location from each species upper thermal limit (upper thermal limit - maximum environmental temperature = warming margin). The latitudinal extents of each functional group were checked prior to analyses to ensure functional group warming margins were comparable, i.e. each functional group had species collected from a broad span of latitudes (Supplementary Figure 1). Due to the lack of thermal limit data collected above 66° latitude, and an uneven sample of functional groups at high latitudes, we limit our global analysis to between the equator and 66° (absolute) latitude (Figure 1, Supplementary Figure 1). Error/variance in upper thermal limit estimates were not reported broadly or consistently across datasets and thus we were unable to perform error propagation analysis throughout the models.

### Global analysis

We examined trends in global species warming margins and how they differ across functional groups and absolute latitude using linear mixed effect models in the lme4 package (Bates et al., 2007). We used a strong inference approach to test and compare three hypotheses: 1) functional group explains most variation in species warming margins, 2) functional group and absolute latitude together explain variation in species warming margins, 3) functional group does not explain variation in species warming margins, but absolute latitude does. To consider variation in the methodology used to estimate species upper thermal limits, methodology was included as a random factor within our models. In addition, to take phylogenetic non-independence into account, we included nested species taxonomic classification (Phylum/ClassOrder/Family/Genus) as a random factor into our models, similar to studies by Lenoir et al. (2020) and Sunday et al. (2011). While it would have been ideal to conduct a full phylogenetic trait analysis, genetic data was not available for many of the species making it impossible to build a robust phylogeny and conduct a reliable comparative phylogenetic analysis. We compared the Akaike Information Criterion (AIC) of each model to determine which hypothesis offered the greatest relative support for understanding the variation in species warming margins as per (Burnham & Anderson, 2002). We also compared each of the three hypotheses with and without the inclusion of method and taxonomic classification as random factors; however, when comparing nested models with different random factor structures, the model with the full random effect structure always best explained variation in species warming margins (Supplementary Table 2).

While it is important to consider phylogenetic non-independence to explain variation in species warming margins and account for taxonomic biases, we also examined trends in warming margins across functional groups without the inclusion of taxonomic non-independence (even though models that included nested taxonomic classification better explained variation in species warming margins). We conducted this additional analysis because closely related species perform similar functional roles within ecosystems (Blondel, 2003) (e.g. all bees are pollinators and most plants are primary producers), and their similar warming margins represent true differences in functional group vulnerabilities. Therefore, by accounting for shared evolutionary history we might be underestimating true differences in functional group vulnerabilities to climate change.

We calculated the proportion of variation in warming margins that each predictor variable and random factor explained using the partR2 package (Stoffel et al., 2021) and the insight package (Lüdecke et al., 2019). We did not include thermogenic capacity (ectothermic animal, endothermic animal, plant and fungi) as a predictor variable because we have already accounted for variation attributed to organism type with nested taxonomic classification as a random factor. However, we appreciate that thermogenic capacity does explain variation in species thermal limits and warming margins as previously described by Bennett et al., (2021) and Sunday et al., (2011 & 2014). Thus, we have outlined differences in thermogenic capacity warming margins for each functional group at the global scale and in low and mid-latitude regions (see Supplementary Information 3), where plant and fungi are independent categories as they use different mechanisms to survive in warm climates than endo- and ectotherms (Angilletta Jr, 2009; Michaletz et al., 2015; Pinkert & Zeuss, 2018). We also appreciate that many groups of species are likely to be under-represented within the dataset, and that biases towards upper thermal limit experiments on certain species could impact our findings. However, we have compared the broadest dataset of terrestrial species upper thermal limits that currently exists in an attempt to analyse the most unbiased and diverse dataset of species warming margins possible. Furthermore, we have included a supplementary file that explicitly lists all of the classes of organisms that contribute towards each functional group, their associated mean warming margins and their count data to outline where biases might occur (Supplementary Information 2). However, by accounting for taxonomic classification as a random factor, we have somewhat accounted for taxonomic biases.

### Regional analysis

While we included absolute latitude as a predictor variable within our global analysis, we wanted to separately examine how functional group vulnerability might change across broad scale regions (tropical vs temperate) with different species compositions. Therefore, to compare functional group vulnerability on a regional scale we split species into low- or mid-latitude regions, where species with collection latitudes between absolute latitudes 0° and 23° were categorised as low-latitude, and species with absolute collection latitudes between 23.1° and 46° were categorised as mid-latitude (the mid-latitude region was limited to 46° due to lack of data over 46° and so that the latitudinal breadths of each geographic region were equal). Using a linear mixed effect model, we examined how functional group vulnerability differed across broad scale geographic regions (low- and mid-latitude) by including an interaction between functional group and region in the model, as well as method and nested taxonomic classification as random factors. To further explore how trends in vulnerability change with changes in species composition across ecosystems we also conducted a local scale case study on the eucalypt forests of south-eastern Australia, the world’s most carbon-dense forests (Keith et al., 2009) (Supplementary Analysis 1). However, due to lack of upper thermal limit data on species across multiple functional groups in other regions, it was not possible to conduct and contrast comparable analyses from other regions.

Significance of effects in linear mixed effect models were assessed using a type II Wald Chi-squared test (in models without interactions) and a type III Wald Chi-squared test in models with interaction terms using the car package (Fox et al., 2012). Method-adjusted functional group means and standard errors were calculated using the emmeans package (Lenth et al., 2018), as were the pairwise contrasts to compare functional group warming margins in low- and mid-latitude regions. Figures were produced using ggplot2 (Wickham, 2011).

### Sensitivity analysis

To determine whether differences between functional group vulnerabilities was underpinned by sample size we conducted a sensitivity analysis. We randomly sampled and bootstrapped upper thermal limit adjusted warming margins from each functional group within the global and regional datasets 10,000 times with sample sizes matching those found in the functional group with the smallest sample size (or region with the lowest sample size for each functional group for the regional analysis) using the sample function in base R. We then calculated functional group warming margin means and 95% confidence intervals from the bootstrapped values and assessed whether they fell within the full dataset 95% confidence intervals (conducted for the global and regional analyses).

## Results

### Global analysis

We found that the model that included functional group, absolute latitude and the random factors upper thermal limit testing methodology and nested taxonomic classification best explained variation in species warming margins across the globe (model *R*^2^ = 0.83) (Supplementary Table 2). Warming margins varied across functional groups (*χ* = 13.18, df = 5, P = 0.0217), where warming margins tended to become narrower with increasing trophic level (except decomposers) (Table 1; Figure 2). Tertiary consumers had the narrowest warming margins and primary producers had the broadest warming margins (Table 1; Figure 2). Pollinators had similar upper thermal limits to primary consumers (as all pollinators in this dataset are also primary consumers), but pollinators had slightly broader warming margins on average. Across all species, absolute latitude played an important role in explaining variation in species warming margins (*χ* = 263.16, df = 1, P < 0.001), where warming margins increased, and thus vulnerability decreased, with latitude (estimate ± SE = 0.19 ± 0.01, t = 16.22, P < 0.001), mirroring the findings of Deutsch et al., (2008) and Sunday et al., (2014).

**Figure 2.**
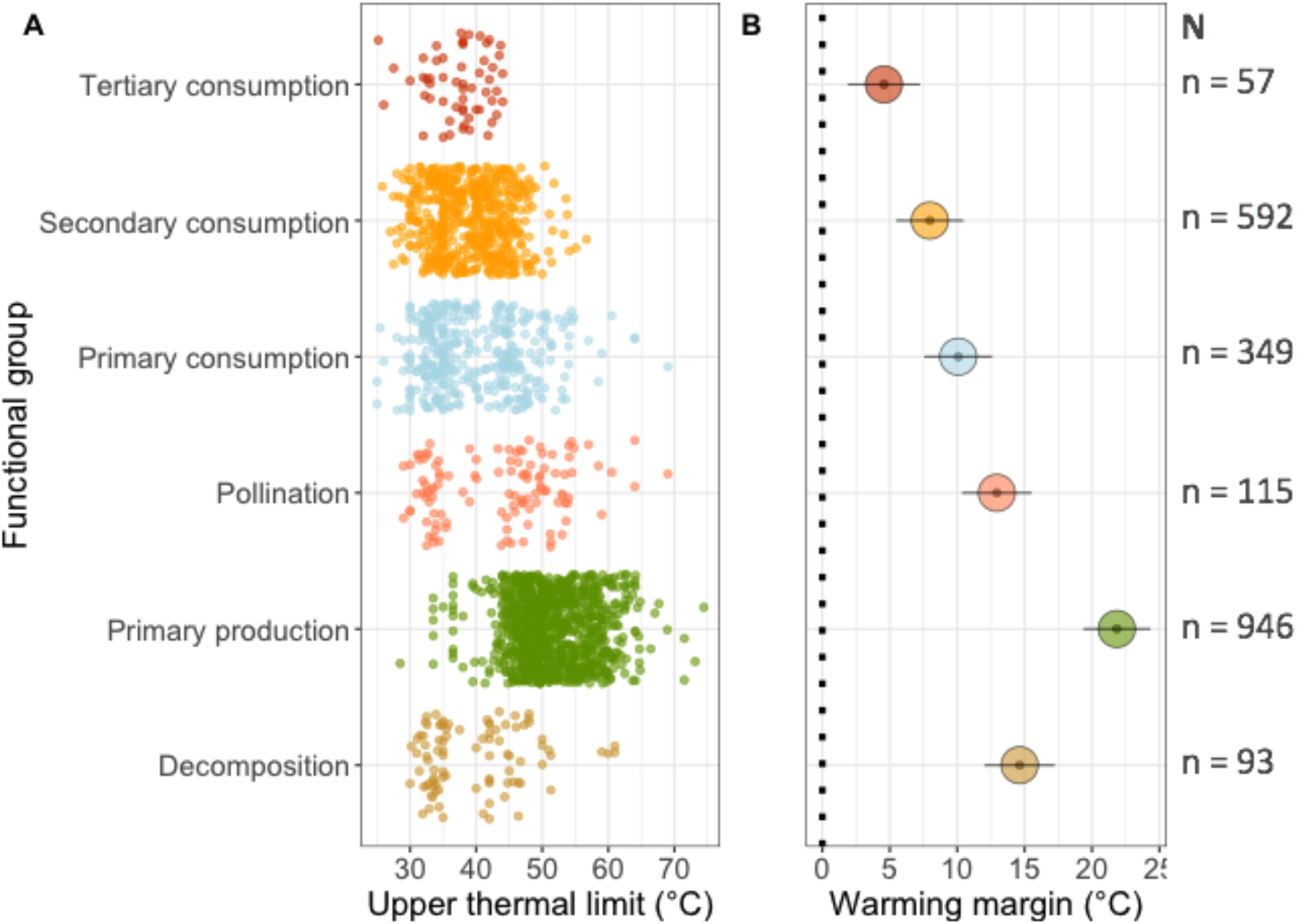
(A) Global comparison of functional group **upper thermal limits** and (B) **warming margin** means and standard error (method adjusted) (degrees Celsius distance between upper thermal limit and maximum environmental temperature (bio5)(Fick & Hijmans, 2017). Sample sizes (N) are located on the right-hand side of the figure.

**Table 1.** Warming margins (estimated model means and standard error calculated from emmeans package) across the globe and in low-latitude (0 - 22° latitude) and mid-latitude regions (22.1 - 44.1° latitude). Means were estimated from a model that included methodology as a random factor.

| **Functional group** | **Global** | **Low-latitude** | **Mid-latitude** |
| --- | --- | --- | --- |
|  | *Mean ± SE* | *Mean ± SE* | *Mean ± SE* |
| Tertiary consumption | 4.37 ± 2.80 | 7.01 ± 3.61 | 4.07 ± 2.89 |
| Secondary consumption | 7.69 ± 2.26 | 7.02 ± 2.72 | 8.28 ± 2.69 |
| Primary consumption | 10.69 ± 2.63 | 11.50 ± 2.74 | 10.41 ± 2.71 |
| Pollination | 12.75 ± 2.70 | 19.23 ± 2.86 | 7.00 ± 2.84 |
| Primary production | 21.85 ± 2.61 | 18.20 ± 2.71 | 22.99 ± 2.68 |
| Decomposition | 14.94 ± 2.74 | 16.05 ± 3.28 | 14.35 ± 2.81 |

While a great deal of the variation in species warming margins was attributed to species evolutionary history (nested taxonomic classification) (67%; Supplementary Table 3), it is important to examine trends in functional group vulnerability without adjusting for shared ancestry as many closely related species often play similar functional roles within ecosystems (Blondel, 2003), representing true variation in functional group warming margins. When evolutionary history is considered, functional group explains 4.0% and absolute latitude accounts for 4.7% of variation in species warming margins, respectively. However, when evolutionary history was not considered, functional group accounted for 36.5% of the variation in species warming margins and absolute latitude accounts for 9.3% of the variation in species warming margins (Supplementary Table 3). We observed the same trend in increasing vulnerability with trophic level from primary producers to tertiary consumers despite whether evolutionary history is considered or not.

The methodology used to estimate upper thermal limit (e.g. dynamic, static or thermal neutral zone) accounted for 8.2% of the variation in species warming margins (Supplementary Table 3). However, the trend of increasing vulnerability with trophic level was not driven by differences in the method used to estimate species upper thermal limits (Supplementary Analysis 2), the presence of nocturnal tertiary consumers (Supplementary Analysis 3), or differences in thermal environments across functional groups (Supplementary Analysis 4). Lastly, we found that global patterns in warming margins across functional groups were robust to sampling bias (Supplementary Table 4).

### Regional analysis

Functional group vulnerability differed between low- and mid-latitude groups (*χ* = 52.22, df = 5, P < 0.001). However, this trend was only driven by differences in warming margins in primary producers across regions, where they were more vulnerable in low-latitude regions than mid-latitude regions (pairwise contrasts between all other functional groups across regions were not significant) (Table 2). Similarly, to the global analysis, there was a general trend in increasing vulnerability (narrower warming margins) with increasing trophic level in both regions (Figure 3). When we excluded evolutionary history (nested taxonomic classification) from our models we found that pollinators were more vulnerable in mid-latitude regions, and that primary producers were more vulnerable in low-latitude regions (Table 2). Regional functional group vulnerability estimates were robust to sampling bias (Supplementary Table 4).

**Figure 3.**
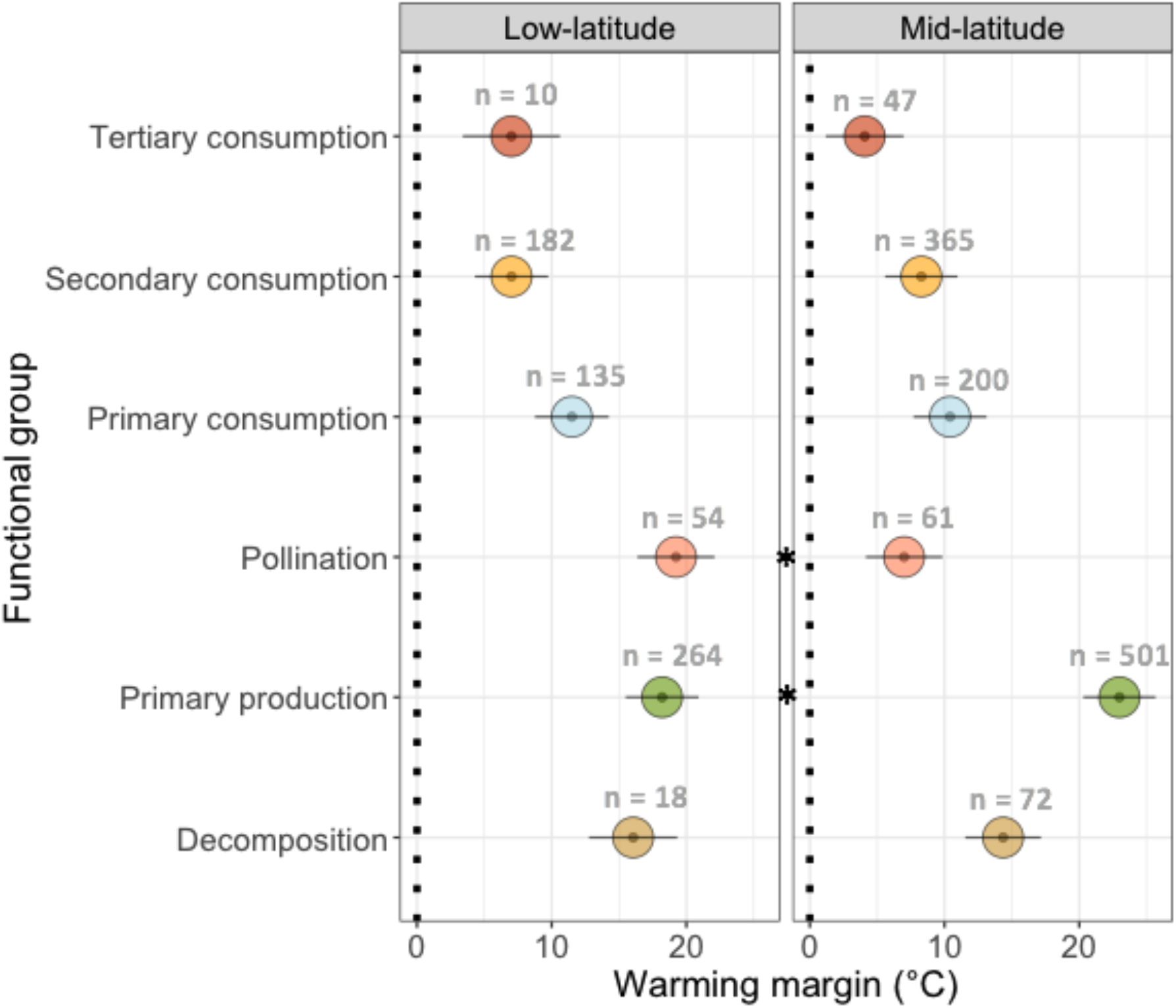
Comparison of low- and mid-latitude functional group warming margin means and standard error (method adjusted). Sample sizes (n) are indicated above each functional group warming margin mean. Asterisks between low- and mid-latitude functional groups indicate significant differences in warming margins between the regions.

**Table 2.** Differences in means of functional group warming margins between low- and mid-latitude regions, with and without the inclusion of evolutionary history (nested taxonomic classification) as a random factor.

| Functional group | Low- vs mid-latitude region pairwise comparisons (considering evolutionary history) |  |  | Low- vs mid-latitude region pairwise comparisons (excluding evolutionary history) |  |  |
| --- | --- | --- | --- | --- | --- | --- |
| | Estimate ± SE | t | P | Estimate ± SE | t | P |
| Tertiary consumption | -1.24 ± 2.59 | -0.48 | 0.629 | -2.94 ± 2.69 | -1.10 | 0.273 |
| Secondary consumption | -0.052 ± 0.57 | -0.09 | 0.928 | 1.27 ± 0.67 | 1.88 | 0.061 |
| Primary consumption | 0.45 ± 0.97 | 0.464 | 0.643 | -1.09 ± 0.85 | 0.85 | 0.199 |
| Pollination | -0.04 ± 1.43 | -0.027 | 0.978 | -12.23 ± 1.45 | 1.45 | < 0.001* |
| Primary production | 5.41 ± 0.58 | 9.40 | < 0.001* | 4.78 ± 0.56 | 8.60 | < 0.001* |
| Decomposition | 1.35 ± 1.99 | 0.69 | 0.493 | -1.70 ± 2.11 | -0.81 | 0.420 |

## Discussion

Understanding how climate change will influence biological systems, from individuals and species to whole ecosystems, is a key goal in ecology (Thakur, 2020; Traill et al., 2010; Tuff et al., 2016; Urban et al., 2016; Zarnetske et al., 2012). A first step towards this objective requires linking species and their physiological vulnerabilities to the functional groups that they contribute to within ecosystems. We compared warming margins of 1,743 terrestrial species across six functional groups (decomposition, primary production, pollination, primary consumption, secondary consumption and tertiary consumption) to determine if functional groups have differential vulnerabilities to climate change, and to assess how unbalanced vulnerabilities might impact ecosystem function. Furthermore, we examined whether certain functional groups are more vulnerable in low-latitude (tropical) regions than mid-latitude (temperate) regions, and if trends in vulnerability are robust to changes in species composition by comparing vulnerabilities of functional groups across geographic scales.

At the global scale, warming margins tended to decrease with trophic level from primary producers (21.85 °C) to tertiary consumers (4.37 °C). This trend was observed when evolutionary history was or was not accounted for. Thus, the true amount of variation in species warming margins that functional group explains is likely to be somewhere in between 4% and 36% (Supplementary Table 3). Because closely related species often play similar functional roles and have similar warming margins, species diversity might only buffer loss of functional roles in ecosystems when species across highly divergent taxonomic groups contribute towards the same functional role. For example, phylum explained the most variation in species warming margins in the nested taxonomic random factor structure (34.5%) (Supplementary Table 3), and thus functional roles in ecosystems that are maintained by highly divergent taxa are likely to have greater warming margin variation, and thus may be more robust to changing climates.

Variation among functional group warming margins appeared to largely reflect variation in upper thermal limits (Figure 2). As species that contributed towards tertiary consumption had the lowest upper thermal limits and the narrowest warming margins, tertiary consumption might be the first functional role to limit ecosystem function across the globe. This finding supports ecological modelling and longitudinal species abundance monitoring studies that suggest vulnerability to climate change will differ across trophic levels, where top predators are likely to be the most at risk (Voigt et al., 2003; Zarnetske et al., 2012; Zhang et al., 2017). Loss of top predators is likely to have prolific knock-on effects throughout ecosystems, where energy and mass cycles become unbalanced (Beschta & Ripple, 2009; Urban et al., 2016; Voigt et al., 2003; Zarnetske et al., 2012; Zhang et al., 2017). For example, the classic reintroduction of tertiary consumers (wolves) into Yellowstone National Park resulted in an increase in woody plants (from controlling the elk population) and an increase in other lower trophic level organism populations (Beschta & Ripple, 2009; Ripple & Beschta, 2012).

While tertiary consumers had the narrowest warming margins overall, at least some species contributing to each consumption group (tertiary, secondary and primary) had warming margins below 0 (Supplementary Figure 2). This indicates that consumer species are already inhabiting environments that experience temperatures higher than their upper thermal limits. It is possible that these species are already using behavioural thermoregulation for survival in their environment (Sunday et al., 2014). Indeed, behavioural thermoregulatory strategies are likely to vary across species, which could also impact trends in functional group vulnerability to climate change. For example, species that use physiological cooling mechanisms (e.g. evaporative cooling) might be less vulnerable to warming climates than species that move to cooler microhabitats (but this is a relatively unexplored topic).

While we can’t explain why we observe a trend of increasing vulnerability with trophic level, our findings align with ecological hypotheses that suggest larger, more active species will be more vulnerable to warming climates due to an inability to maintain energetic requirements in hot conditions (metabolic meltdown) (Brown et al., 2004; Enquist et al., 2003; Huey & Kingsolver, 2019; Vasseur & McCann, 2005; Voigt et al., 2003). This idea is supported by a study that examined trends in upper thermal limits and body mass in grasshoppers, where the largest individuals had the lowest upper thermal limits (Youngblood et al., 2019), however, in other systems, body mass does not explain variation in upper thermal limits, e.g. Fijian bees (da Silva et al., 2021). We were unable to examine how body mass correlates with warming margins and functional groups across geographic space due to a lack of publicly available body mass data for many of the species in our dataset. However, an analysis of this kind will be important in future studies and we encourage the inclusion of body mass in publicly available datasets in future.

Primary producers had very high upper thermal limits and broad warming margins on average compared to all other functional groups (Figure 2, Table 2). Primary producers might have evolved high upper thermal limits as a mechanism to survive in fluctuating environments as plants are usually stationary, and thus require the capacity to acclimate with thermal change or maintain broad thermal tolerances to survive with changes in environmental temperature (Huey et al., 2003). High thermal tolerances have also been observed in stationary developmental life stages of species such as *Drosophila* (Moghadam et al., 2019), supporting this idea. Within each functional group we observed a great deal of variation in thermal limits and warming margins (Supplementary Figure 2). Variation in upper thermal limits between species might allow key functional roles to be maintained within ecosystems as backup species with high upper thermal limits can perpetuate functional roles if species with lower upper thermal limits are lost (i.e. functional redundancy). However, global functional redundancy has little relevance for local redundancy (i.e. a species that provides a certain functional role will not help maintain ecosystem function in an environment where that species is lost), and thus it is important to assess whether climate change vulnerability differs across regions, as species composition changes for different functional groups.

We found that trends in functional group vulnerability within each broad-scale region mirrored trends in vulnerability at the global scale (increasing vulnerability with increasing trophic level). However, there was little variation between tertiary consumer and secondary consumer warming margins in the low-latitude region, potentially due to geographic sampling biases, where wet-tropical regions and developing nations tend to be under sampled compared to temperate regions (White et al., 2021) (Figure 1; Supplementary Figure 1). In the global analysis species warming margins increased with absolute latitude, suggesting that species that inhabit low-latitude regions are more vulnerable to climate change. However, when we split low- and mid-latitude regions into two broad scale regions we found that primary producers are more vulnerable in low-latitude regions and pollinators are likely to be more vulnerable in mid-latitude regions (Table 2). Thus, vulnerability to climate change is likely to depend on the thermal environment species inhabit (e.g. tropical or temperate), the functional role they play in ecosystems, and their evolutionary histories. Furthermore, in the local south-eastern Australia case study, we found that tertiary consumers were more resilient than other functional groups (Supplementary Analysis 1), suggesting that functional group vulnerability trends might not be unanimous across ecosystems composed of different species. However, more data are required to make improved inferences on how functional group vulnerability changes across ecosystems and species compositions.

### Study limitations

The main limitation of this study is not being able to account for plasticity or evolutionary potential in our analyses (Supplementary Information Section 4). Species are likely to be able to shift their thermal limits via plasticity or evolution, but a lack of data on the extent to which plasticity and evolutionary potential across species limits our capacity to factor evolution into vulnerability estimates. Nevertheless, the extent to which evolution is likely to shift upper thermal limits is likely to be small and unlikely to match the pace of climate warming (Gunderson & Stillman, 2015; Kellermann & van Heerwaarden, 2019). We also acknowledge that upper thermal limits are likely to under-estimate climate change vulnerability because rising temperatures will impose a range of sublethal effects on fitness at temperatures below species upper thermal limits (da Silva et al., 2020; van Heerwaarden & Sgrò, 2021). So while the absolute values of warming margins are unlikely to reflect the exact temperatures at which species will be negatively impacted by climate change, warming margins are still likely to capture an element of fitness/sub-lethal effects on phenotypes and hence the rank order of climate change vulnerability will remain (van Heerwaarden & Sgrò, 2021). Further research that seeks to examine plastic responses, evolutionary potential and sub-lethal effects of temperature on species abilities to perform functional roles will be important in improving estimates of functional group vulnerability. In addition, considering how variation in climate velocity (i.e. rate of climate change) across space is needed as not all regions are warming at the same rate (VanDerWal et al., 2013). Thus, species that have the same warming margins but inhabit regions with different climate velocity will differ in their vulnerability.

Finally, many groups of species are underrepresented in this analysis, we examined the warming margins of 1,743 species which is well below the number of species that exist on the planet (estimates suggest between 5.3 million – 1 trillion (Locey & Lennon, 2016)) and contribute towards functional roles in the ecosystem. Thus, further studies that continue to estimate species thermal tolerances will be important for gaining a more robust understanding of how functional group vulnerabilities vary across the globe and local scale ecosystems.

## Conclusions

Our results demonstrate how global drivers of biodiversity loss (climate change) are non-random with respect to the function roles that species play in ecosystems. Species that contribute to higher trophic levels are likely to be the most vulnerable to further climate warming and primary producers are likely to be the most resilient to increases in environmental temperature. This trend of increasing vulnerability to climate change with trophic level is observed at the global scale as well as within broad scale tropical and temperate regions. Importantly, however, vulnerability trends might differ with changes in species composition across fine scale ecosystems, and thus caution should be used when transferring global vulnerability trends to local ecosystems. Overall, we observed variation in the climate change vulnerability of functional groups across all geographic scales, which is likely to disrupt functional group interactions and eventually impact ecosystem function (Voigt et al., 2003). Thus, governments and private organisations should seek to reduce carbon emissions to conserve ecosystem function for the future.

