## Supplementary information for "Linking physiology to ecosystem function: how vulnerable are different functional groups to climate change?"

**Supplementary material section 1.**

**Search criteria for pollinators, decomposers and tertiary consumers**

Addition of thermal tolerance data to GlobTherm (Bennett et al. 2018) and Lancaster and Humphreys (2020) datasets. Searched on the Web of Science website. Data must be from terrestrial field collected organisms (no lab strains or cultures), and must have a precise collection latitude and longitude.

**Search terms for pollinators**

(pollinator, CTmax, upper thermal limit)

(pollinator, CTmax, upper thermal limit, insect)

(pollinator, CTmax, upper thermal limit, bird)

**List of citations added to the database**

Warren et al. 2010 (fig wasps)

Oyen and Dillon 2016 (bees)

Hamblin et al. 2017 (bees)

Burdine and McCluney 2019 (bees)

Gonzalez et al. 2020 (bees)

Silva et al. 2020 (butterfly)

Dongmo et al. 2021 (butterfly)

da Silva et al. 2021(bees)

**Search terms for decomposers:**

(Decomposer, CTmax, upper thermal limit)

(Fungi, maximum growth temperature)

(Decomposer, CTmax, upper thermal limit, insect)

Nervo et al. 2021 (dung beetles)

Eggins & Coursey 1964 (fungi)

Rinu et al 2011 (fungi)

Harris and Wright 2021 (Isopod)

Janowiecki et al. 2019 (termites)

**Search terms for tertiary consumer:**

(Tertiary consumer, upper thermal limit, thermal tolerance)

(Top predator, upper thermal limit, thermal tolerance)

(Carnivor, upper thermal limit, thermal tolerance)

No new data on tertiary consumer upper thermal limits

### Supplementary Material Section 2

#### Supplementary figures and tables

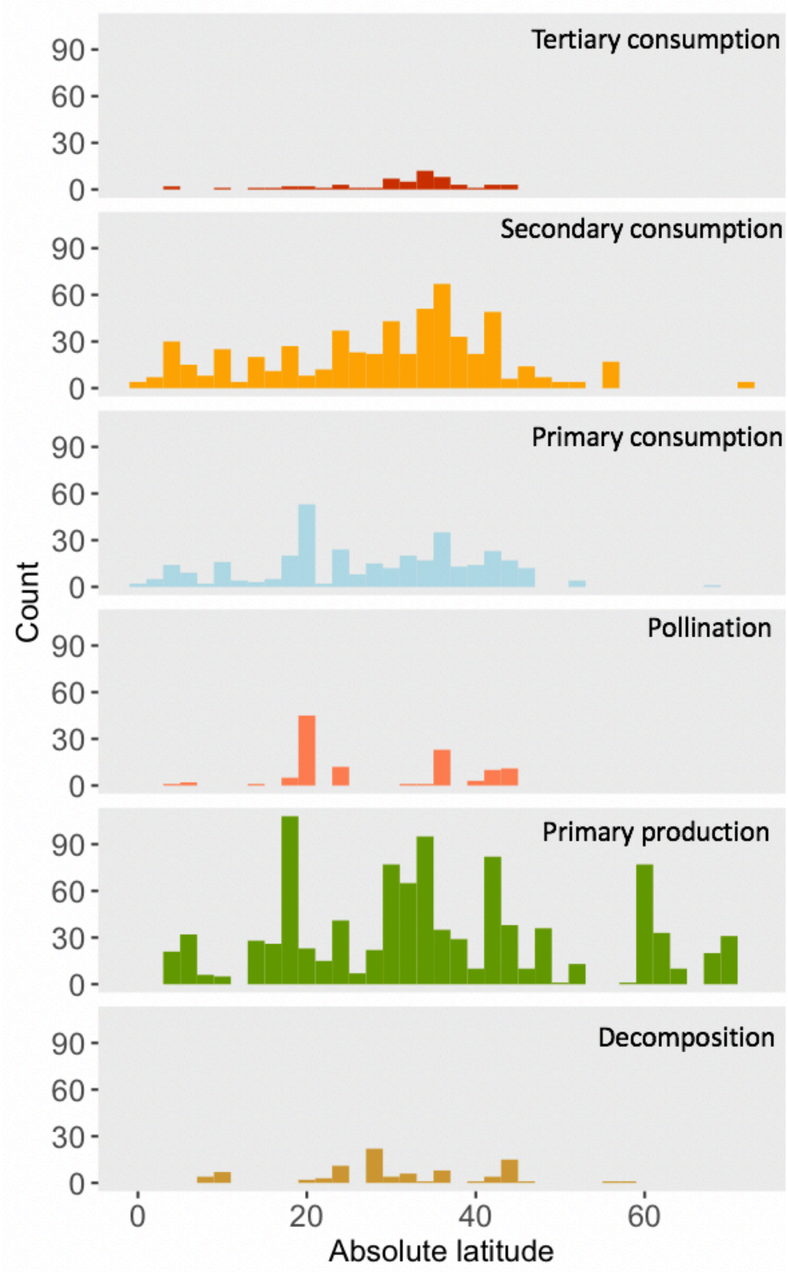


**Supplementary Figure 1.** Absolute latitudinal spread of thermal tolerance data for each functional group.

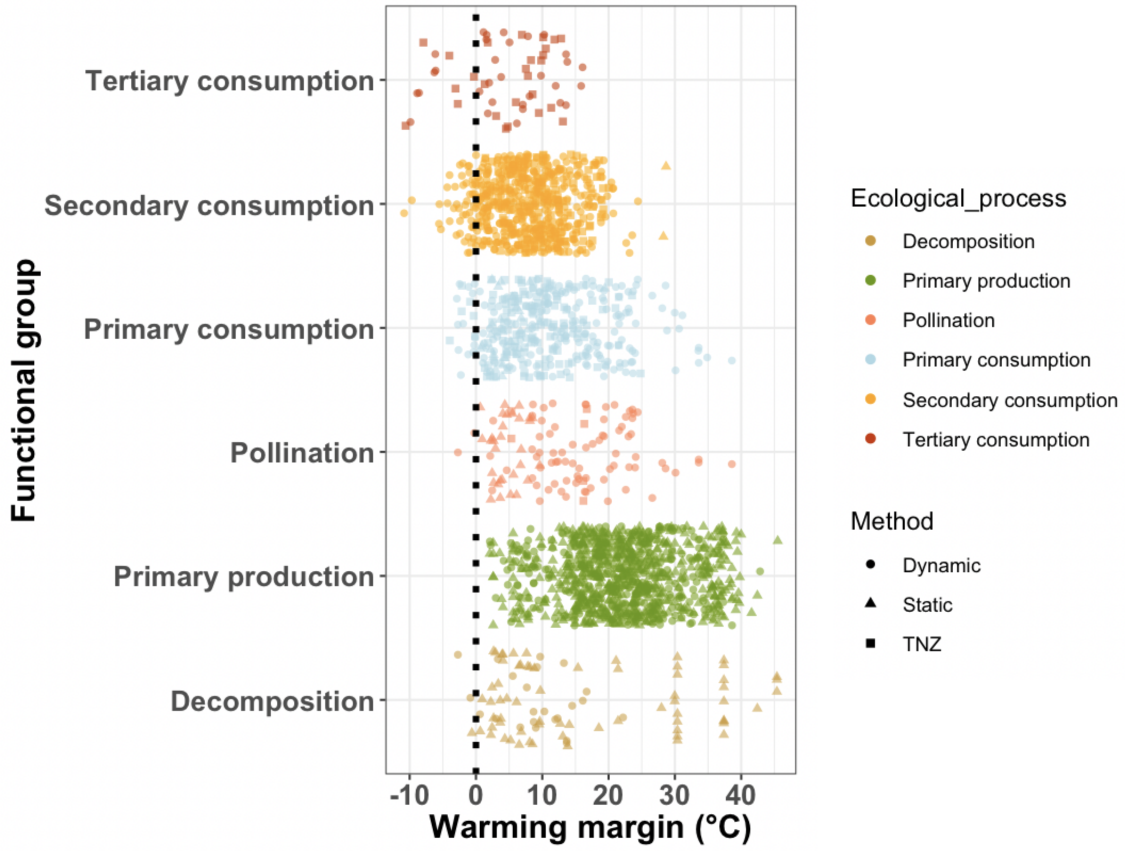


**Supplementary Figure 2.** Global functional group warming margins (method adjusted) (degrees difference between upper thermal limit and average maximum environmental temperature (Bio5).

**Supplementary Table 1.** Summary of methodology categories used to estimate upper thermal limits of species that contribute to each functional group. The class of organism measured, the number of species/populations measured and the warming margin is also provided. Within the dynamic category studies use a range of ramping rates and dynamic metrics. Within the static method categories studies use LT50, LT100, and studies on fungi in particular assess the maximum temperature at which they will grow. Finally, the upper edge of the thermal neutral zone (TNZ) is used to assess the upper limit of many bird and mammal species, and yields lower upper thermal limit estimates than static or dynamic methods on average. However, when studies that use TNZ are removed from the analysis the main conclusions we draw do not change (Supplementary Material Section 3, Analysis 2).

| Functional group and experimental method | Class | Count | Mean warming margin |
|---|---|---|---|
| **Tertiary consumption** | | **57** | **4.61** |
| Dynamic | Lepidosauria | 31 | 8.38 |
| TNZ | Aves | 11 | 0.89 |
| | Mammalia | 15 | -0.44 |
| **Secondary consumption** | | **596** | **9.60** |
| Dynamic | Amphibia | 108 | 6.18 |
| | Arachnida | 20 | 21.28 |
| | Archelosauria | 2 | 10.23 |
| | Collembola | 3 | 34.12 |
| | Insecta | 139 | 13.75 |
| | Lepidosauria | 152 | 11.52 |
| Static | Arachnida | 2 | 28.47 |
| | Lepidosauria | 1 | 6.05 |
| TNZ | Aves | 58 | 5.46 |
| | Mammalia | 111 | 4.16 |
| **Primary consumption** | | **350** | **11.48** |
| Dynamic | Insecta | 100 | 21.11 |
| | Lepidosauria | 92 | 13.23 |
| Static | Gastropoda | 1 | 25.60 |
| | Insecta | 32 | 5.17 |
| TNZ | Aves | 35 | 6.98 |
| | Mammalia | 90 | 2.82 |
| **Pollination** | | **115** | **15.52** |
| Dynamic | Insecta | 78 | 20.41 |
| Static | Insecta | 31 | 4.71 |

| | | | |
|---|---|---|---|
| TNZ | Aves | 6 | 7.77 |
| **Primary production** | | **997** | **24.49** |
| Dynamic | Ginkgoopsida | 1 | 33.09 |
| | Lecanoromycetes | 1 | 26.20 |
| | Magnoliopsida | 377 | 25.97 |
| | Pinopsida | 23 | 27.54 |
| | Polytrichopsida | 1 | 24.63 |
| | Spermatophyta | 2 | 27.87 |
| Static | Bryophyta | 1 | 5.60 |
| | Bryopsida | 4 | 17.23 |
| | Jungermanniopsida | 18 | 14.49 |
| | Magnoliopsida | 521 | 23.53 |
| | Pinopsida | 15 | 30.40 |
| | Polypodiopsida | 27 | 21.29 |
| | Sphagnopsida | 6 | 37.90 |
| **Decomposition** | | **91** | **16.25** |
| Dynamic | Arthropoda | 3 | 8.49 |
| | Insecta | 16 | 13.33 |
| | Lecanoromycetes | 3 | 16.54 |
| | Malacostraca | 1 | 17.28 |
| Static | Chytridiomycetes | 2 | 9.16 |
| | Euascomycetes | 2 | 36.20 |
| | Eurotiomycetes | 22 | 33.76 |
| | Insecta | 41 | 7.55 |
| | Sordariomycetes | 1 | 30.38 |

**Supplementary Table 2**. Model comparison using AIC to determine which random factor structure offered the greatest relative support for understanding the variation in species warming within each hypothesis. Models that included both methodology and taxonomic classification had the lowest AICs. Thus we also compared model support across hypotheses/models that included the full random effect structure.

| Model | Random effect structure | AIC | Within hyp. Δ AIC | Across hyp. Δ AIC |
|---|---|---|---|---|
| **H1. Functional group** | Methodology + Taxonomic classification | 13791.17* | 0 | 230.49 |
| | Taxonomic classification | 13898.89 | 107.72 | |
| | Methodology | 15019.06 | 1227.89 | |
| | No random factors | 15329.86 | 1538.69 | |
| **H2. Functional group + absolute latitude** | **Methodology + Taxonomic classification** | **13560.68**** | **0** | **0** |
| | Taxonomic classification | 13729.08 | 168.4 | |
| | Methodology | 14803.68 | 1243 | |
| | No random factors | 15153.46 | 1592.78 | |
| **H3. Absolute latitude** | Methodology + Taxonomic classification | 13573.2* | 0 | 12.52 |
| | Taxonomic classification | 13740.15 | 166.95 | |
| | Methodology | 15598.99 | 2025.79 | |
| | No random factors | 16130.59 | 2557.39 | |

**Supplementary Table 3.** Variance and percent variance of species warming margin data that each fixed effect and random effect explains within the global warming margin analysis. The model included functional group and latitude as fixed effects and the method used to estimate upper thermal limits and nested taxonomic classification (Phylum/Class/Order/Family/Genus) as random effects. The model had a total $R^2$ of 0.833.

| **Model including methodology and nested taxonomic classification as random factors** | | |
|---|---|---|
| Fixed and random effects | Variance | % variance |
| Fixed effects (Functional group + absolute latitude) | 12.04 | 9.07 |
| Random: Phylum/Class/Order/Family/Genus | 12.228 | 9.21 |
| Random: Phylum/Class/Order/Family | 3.764 | 2.84 |
| Random: Phylum/Class/Order | 5.884 | 4.43 |
| Random: Phylum/Class | 32.265 | 24.31 |
| Random: Phylum | 34.999 | 26.37 |
| Random: Method | 10.87 | 8.19 |
| Residuals | 20.65 | 15.56 |
| **Total variance** | **132.7** | **100.00** |
| **Total variation attributed to nested taxonomic classification** | **89.14** | **67.17** |
| **Unique fixed effect variation (estimated from marginal $R^2$ values (partR2 package)).** *Note functional group and absolute latitude uniquely explains 4.0% and 4.7% respectively, of the variation in species warming margins. However, together they only explain 12.04%. This is typical when predictors are somewhat correlated and some of the variance is explained by both predictors (Stoffel et al., 2021) | | |
| **Functional group** | | **4.0** |
| Absolute latitude | | 4.7 |
| **Model excluding taxonomic classification as a random factor** | | |
| Fixed effects (Functional group + absolute latitude) | 53.55 | 41.8 |
| Random: Method | 18.02 | 14.1 |
| Residual | 56.59 | 44.2 |
| **Total variance** | **128.16** | **100** |
| **Unique fixed effect variation** | | |
| **Functional group** | | **36.5** |
| Absolute latitude | | 9.26 |

**Supplementary Table 4.** Global and regional sensitivity analyses to examine whether trends in functional group vulnerability are robust. We used the method adjusted warming margin values for each species so that means and confidence limits were comparable to the model means. Values were randomly sampled bootstrapped 10,000 times.

| **Functional group** | **Random sample bootstrap mean** | **Random sample bootstrap lower 95% CL** | **Random sample bootstrap Upper 95% CL** |
|---|---|---|---|
| *Global Analysis Bootstrap Results* | | | |
| **Tertiary Consumers** | 4.66 | 3.13 | 6.16 |
| **Secondary Consumers** | 7.99 | 6.72 | 9.30 |
| **Primary Consumers** | 10.98 | 9.31 | 12.69 |
| **Pollinators** | 13.05 | 11.21 | 14.88 |
| **Primary Producers** | 22.15 | 21.71 | 22.60 |
| **Decomposers** | 15.22 | 12.98 | 17.48 |
| *Low Latitude Analysis Bootstrap Results* | | | |
| **Tertiary Consumers** | 7.05 | 5.08 | 9.05 |
| **Secondary Consumers** | 7.04 | 6.39 | 7.71 |
| **Primary Consumers** | 11.60 | 10.41 | 12.81 |
| **Pollinators** | 19.35 | 17.81 | 20.89 |
| **Primary Producers** | 18.70 | 17.95 | 19.47 |
| **Decomposers** | 16.53 | 13.67 | 19.55 |
| *Mid latitude Analysis Bootstrap Results* | | | |
| **Tertiary Consumers** | 4.18 | 0.32 | 7.95 |
| **Secondary Consumers** | 7.52 | 6.84 | 8.18 |
| **Primary Consumers** | 9.68 | 8.74 | 10.64 |
| **Pollinators** | 7.44 | 6.39 | 8.50 |
| **Primary Producers** | 20.41 | 19.71 | 21.09 |
| **Decomposers** | 14.46 | 9.24 | 20.05 |

### Supplementary material section 3

#### Supplementary Analysis 1

#### Local scale analysis methods

To explore whether trends in functional group vulnerability are maintained across geographic scale (global - regional - local) and with changes in species composition, we conducted a local scale analysis comparing warming margins across functional groups in south-eastern Australian eucalypt forests. This ecosystem was chosen for the local scale analysis due to the high sampling of species upper thermal limits (N = 136) compared to other local ecosystems across the globe. For this analysis, species that inhabit latitudes between -26° to -39° and longitudes between 142° to 160° were analysed (desert species with collection latitudes inland from the Great Dividing Range were eliminated as they persist in a very different bioregion to the rest of southeastern Australia). A linear mixed model was used to assess how warming margins differ across functional groups at a more localised scale. To assess if differences in the vulnerability of functional groups in southeastern Australia were driven by small sample sizes or if they truly reflected local scale differences in functional group vulnerability, we randomly sampled and bootstrapped warming margins from the global dataset 10,000 times with sample sizes matching those found in the southeastern Australian dataset. We then calculated means and 95% confidence intervals from the global bootstrapped values and assessed whether the southeastern Australian functional group means fell within the global bootstrapped global 95% confidence intervals.

#### Local scale analysis results

The south-eastern Australian eucalypt forests are the world's most carbon-dense forests (Keith et al., 2009), and they are the only region in the world where data for over 100 species within a particular environment type exists, making it the best case study to assess ecosystem vulnerability at a more localised scale. Again, warming margins varied across functional groups ($\chi_3$ = 14.5, P = 0.002; Supplementary Table 7). However, tertiary consumers tended to have broader warming margins compared to the rest of the globe (Supplementary Table 7; Supplementary Table 3), but the vulnerability of all other functional groups was not different to randomly sampled global functional group vulnerabilities (Supplementary Figure 3). There was no available data on pollinator thermal tolerance in south-eastern Australia and only 1 data point was available for decomposers which was excluded from the south-eastern Australian analyses.

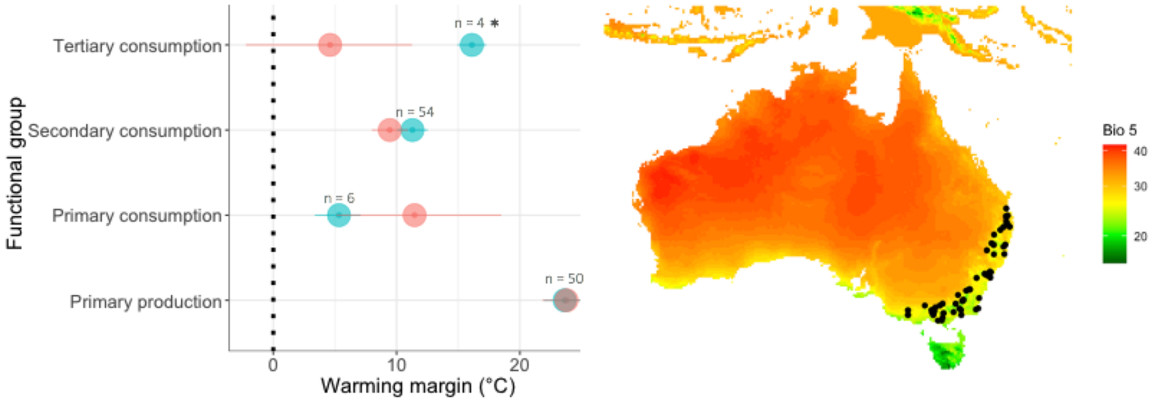


**Supplementary Figure 3**. Means and 95% confidence intervals of southeastern Australian functional group warming margins (blue) and randomly sampled (sample sizes matched those available for southeastern Australia) bootstrapped global warming margin means and 95% confidence intervals (red). Asterisks indicate where south-eastern Australian warming margins were significantly different from global functional group warming margins. There was no decomposer or pollinator thermal tolerance data for this region. The right-hand panel shows the southeastern Australian eucalypt forest region we refer to and the BioClim data Bio 5 across the country (mean maximum temperatures from the hottest month).

**Supplementary 7.** Bootstrapped warming margin means and 95% upper and lower confidence limits of the south-east Australian dataset and the global warming margins dataset where sample sizes matched those in the south-eastern Australia dataset. Only one data point was available for decomposers, so it was eliminated from the analysis and no pollinator data was available.

| **Functional group** | ***Mean*** | ***Lower confidence limit*** | ***Upper confidence limit*** | ***Location*** |
|---|---|---|---|---|
| Tertiary consumption | 16.11 | 14.94 | 17.27 | SE Australia |
| Secondary consumption | 11.29 | 10.06 | 12.53 | SE Australia |
| Primary consumption | 5.33 | 3.38 | 7.09 | SE Australia |
| Primary production | 23.63 | 21.93 | 25.35 | SE Australia |
| Tertiary consumption | 4.61 | -2.22 | 11.25 | Global bootstrap |
| Secondary consumption | 9.45 | 7.99 | 10.94 | Global bootstrap |
| Primary consumption | 11.46 | 5.11 | 18.49 | Global bootstrap |
| Primary production | 23.78 | 21.84 | 25.73 | Global bootstrap |

**Local scale analysis discussion**

Within the south-eastern Australian eucalypt forests case study, we found that tertiary consumers had broader warming margins compared to tertiary consumers randomly sampled from the global dataset. The tendency for higher warming margins in tertiary consumers could be because snakes made up all the data in that functional group in south-eastern Australia, while the global sample comprised mammals, birds and reptiles. While some other terrestrial tertiary consumers are found in south-eastern Australia (e.g. owls, eagles, feral cats and dogs), there are very few top-predators found in this environment. Many ecosystems, such as islands, may have limited diversity within certain functional groups, which could influence local scale vulnerability patterns. For example, after the accidental introduction of the brown tree snake in Guam, bird, bat and lizard populations plummeted (Fritts & Rodda, 1998). Thus, in consumption and pollination processes will be limited towards taxa such as insects, snakes, and some small mammal species, potentially skewing the vulnerability of those functional groups at the local scale. South-eastern Australian eucalypt forests represent the most highly sampled (for upper thermal limit data) ecosystem in the world, yet we still have limited power to confidently make inferences regarding which functional roles are likely to limit ecosystem function with further climate change. Thus, at this stage, we can only recommend for caution to be used when transferring global vulnerability trends to local scales. We recommend for future research to compare local scale functional group vulnerabilities at multiple locations across the globe to gain a better understanding of vulnerability trends across space and species compositions.

*Supplementary analysis 2*

Global analysis excluding TNZ (upper limit of thermal neutral zone) collected upper thermal limit data.

The exclusion of TNZ does not change our findings that tertiary consumers have the narrowest warming margins (Supplementary Table 7).

We used a strong inference approach to compare three hypotheses (excluding TNZ collected data).

H1. Warming margin ~ functional group + absolute latitude + (1|method) + (1|Phylum/Class/Order/Family/Genus).

AIC = 11557.32 *

H2. Warming margin ~ functional group + (1|method) + (1|Phylum/Class/Order/Family/Genus).

AIC = 11771.61

H3. Warming margin ~ absolute latitude + (1|method) + (1|Phylum/Class/Order/Family/Genus).

AIC = 11565.35

**Table 8**. Estimated model mean warming margins of global upper thermal limit data without TNZ data generated from H1 model.

| Functional group | Est. marginal mean warming margin | SE |
|---|---|---|
| Tertiary consumption | 12.1 | 4.61 |
| Secondary consumption | 15.0 | 4.34 |
| Primary consumption | 17.0 | 4.18 |
| Pollination | 17.1 | 4.19 |
| Primary production | 19.3 | 4.40 |
| Decomposition | 17.3 | 4.13 |

*Supplementary analysis 3*

**Elimination of nocturnal tertiary consumers**

Elimination of nocturnal tertiary consumers (owls and bats) does not impact the finding that tertiary consumers have the narrowest warming margin (Supplementary Table 8).

We used a strong inference approach to compare three hypotheses (excluding nocturnal animals).

H1. Warming margin ~ functional group + absolute latitude + (1|method) + (1|Phylum/Class/Order/Family/Genus).

AIC = 13485.76*

H2. Warming margin ~ functional group + (1|method) + (1|Phylum/Class/Order/Family/Genus).

AIC = 13716.43

H3. Warming margin ~ absolute latitude + (1|method) + (1|Phylum/Class/Order/Family/Genus).

AIC = 13496.49

**Supplementary Table 9.** Estimated marginal mean warming margins of global upper thermal limit data without nocturnal tertiary consumers.

| Functional group | Est. marginal mean warming margin | SE |
|---|---|---|
| Tertiary consumption | 11.3 | 4.13 |
| Secondary consumption | 14.6 | 3.92 |
| Primary consumption | 16.0 | 3.87 |
| Pollination | 16.1 | 3.88 |
| Primary production | 18.7 | 4.23 |
| Decomposition | 16.4 | 3.84 |

*Supplementary analysis 4*

**Do species that contribute towards different functional groups inhabit different maximum thermal environments?**

**Model structure**

Mean maximum temp of hottest month ~ functional group + (1| Phylum/Class/Order/Family/Genus)

**Anova results**

Species that contribute towards different functional roles do not inhabit different mean thermal environments ($F = 0.8036$, $df = 5$, $P = 0.552$).

**Supplementary Information Section 4**

**Research priorities:**

*Prioritise research on species that contribute importantly to functional roles, but which currently have small sample sizes.*

While it will only ever be possible to test the thermal limits of a sample of the species on earth, it is important to obtain a representative sample of species that contribute to different functional roles across different environments. For example, relatively little data exists on the thermal limits of tertiary consumers, pollinators and decomposers across most regions of the globe. A great diversity of species contribute to these processes and therefore we currently have a rudimentary understanding of how they will respond to climate change across diverse ecosystems.

*Adding evolution and plasticity into vulnerability estimates*

Based on current efforts to catalogue species vulnerabilities to climate change we know that upper thermal limits vary within and across species (Bennett et al., 2021; Castañeda et al., 2015; da Silva et al., 2021; Diamond, 2018; Kellermann et al., 2012; Kingsolver et al., 2013). We made the simplifying assumption that evolution won't shift upper thermal limits because we currently have insufficient interspecific data on evolutionary potential and plasticity. Predicting the degree to which evolution will shift upper thermal limits requires carefully controlled experiments to untangle environmental and genetic effects on phenotypes (Hoffmann & Sgro, 2018). This is difficult for many species, but doing so in the species we can, could help build a general understanding of the potential for evolution to shift upper thermal limits. Our understanding on the role of plasticity in upper thermal limits (i.e. heat hardening and acclimation capacity) is also growing but understanding the relative importance of plasticity also requires carefully controlled experiments (Gunderson & Stillman, 2015; Kellermann et al., 2020). Determining the degree to which thermal limits are likely to shift via plasticity and evolution will likely improve our ability to predict which functional groups are most at risk and likely to limit ecosystem function in the future.

Although this is an ambitious research agenda, establishing linkages between species climate vulnerabilities (which include evolutionary potential and plasticity estimates) and ecosystem function will ideally enable more realistic forecasts of responses to climate change and thus promote informed conservation planning and management decisions.

**Supplementary material references**

Bennett, J. M., Sunday, J., Calosi, P., Villalobos, F., Martínez, B., Molina-Venegas, R., Araújo, M. B., Algar, A. C., Clusella-Trullas, S., & Hawkins, B. A. (2021). The evolution of critical thermal limits of life on Earth. *Nature Communications*, *12*(1), 1–9.

Castañeda, L. E., Rezende, E. L., & Santos, M. (2015). Heat tolerance in *Drosophila subobscura* along a latitudinal gradient: Contrasting patterns between plastic and genetic responses: Plastic and genetic responses of heat tolerance. *Evolution*, *69*(10), 2721–2734. https://doi.org/10.1111/evo.12757

da Silva, C. R., Beaman, J. E., Dorey, J. B., Barker, S. J., Congedi, N. C., Elmer, M. C., Galvin, S., Tuiwawa, M., Stevens, M. I., & Alton, L. A. (2021). Climate change and invasive species: A physiological performance comparison of invasive and endemic bees in Fiji. *Journal of Experimental Biology*, *224*(1).

Diamond, S. E. (2018). Contemporary climate-driven range shifts: Putting evolution back on the table. *Functional Ecology*, *32*(7), 1652–1665. https://doi.org/10.1111/1365-2435.13095

Fritts, T. H., & Rodda, G. H. (1998). The role of introduced species in the degradation of island ecosystems: A case history of Guam. *Annual Review of Ecology and Systematics*, *29*(1), 113–140.

Gunderson, A. R., & Stillman, J. H. (2015). Plasticity in thermal tolerance has limited potential to buffer ectotherms from global warming. *Proceedings of the Royal Society B: Biological Sciences*, *282*(1808), 20150401.

Hoffmann, A. A., & Sgro, C. M. (2018). Comparative studies of critical physiological limits and vulnerability to environmental extremes in small ectotherms: How much environmental control is needed? *Integrative Zoology*, *13*(4), 355–371.

Keith, H., Mackey, B. G., & Lindenmayer, D. B. (2009). Re-evaluation of forest biomass carbon stocks and lessons from the world's most carbon-dense forests. *Proceedings of the National Academy of Sciences*, *106*(28), 11635–11640.

Kellermann, V., McEvey, S. F., Sgrò, C. M., & Hoffmann, A. A. (2020). Phenotypic Plasticity for Desiccation Resistance, Climate Change, and Future Species Distributions: Will Plasticity Have Much Impact? *The American Naturalist*, *196*(3), 306–315.

Kellermann, V., Overgaard, J., Hoffmann, A. A., Fløjgaard, C., Svenning, J.-C., & Loeschcke, V. (2012). Upper thermal limits of Drosophila are linked to species distributions and strongly constrained phylogenetically. *Proceedings of the National Academy of Sciences*, *109*(40), 16228–16233.

Kingsolver, J. G., Diamond, S. E., & Buckley, L. B. (2013). Heat stress and the fitness consequences of climate change for terrestrial ectotherms. *Functional Ecology*, *27*(6), 1415–1423.

Stoffel, M. A., Nakagawa, S., & Schielzeth, H. (2021). partR2: Partitioning R2 in generalized linear mixed models. *PeerJ*, *9*, e11414.
